# Antioxidants maintain Butyrate production by Human Gut *Clostridia* in the presence of Oxygen

**DOI:** 10.1101/652958

**Authors:** Matthieu Million, Nicholas Armstrong, Saber Khelaifia, Elodie Guilhot, Magali Richez, Jean-Christophe Lagier, Gregory Dubourg, Eric Chabriere, Didier Raoult

## Abstract

**Background:** Oxygen diffused from the human gut mucosa and shape the microbiota with a radial gradient of microbes according to their oxygen tolerance, while microbial and chemical oxygen consumption maintains the lumen in a deeply anaerobic state. Uncontrolled oxidative stress and hyperoxygenation have been reported as a pathogenic mechanism in *Salmonella* or *Citrobacter rodentium* infection, in patients with HIV and in severe acute malnutrition. We recently found that antioxidants allow strict anaerobes, including methanogenic archaea, to thrive in an oxidative environment (aerobic). Here, we tested the metabolomics switching of the 3 most odorous anaerobic microbes isolated from human gut when grown in aerobiosis with antioxidants.

**Methods:** Three human gut Clostridia, *Clostridium sporogenes, Clostridium lituseburense* and *Clostridium subterminale*, isolated by culturomics, were grown in anaerobiosis or in aerobiosis with antioxidants. Gaz and liquid chromatography-Mass spectrometry (GC/MS and LC/MS) were used for metabolomics analysis.

**Results:** An unexpected global dichotomic metabolomic switching from thiols, alcohols and short-chain fatty acid esters to a specific aerobic metabolic repertoire with the production of alkanes, cycloheptatriene and, paradoxically, increased butyrate production, was observed. Analysis of polar metabolites confirmed the discovery of an unexplored aerobic metabolic repertoire, including the production of specific dipeptides and several lysophospholipids, thus unraveling unsuspected human gut microbiome capacities.

**Conclusions:** Antioxidants unraveled an unexplored aerobic metabolic repertoire of human gut *Clostridia*. The increased production of butyrate suggests that antioxidants contribute to the maintenance and the active resilience of the human gut microbiome against oxidative aggression, as during *Salmonella* infection.

## BACKGROUND

Odorants are volatile organic compounds detected by the human nose that generate an olfactive signal mainly comprising thiols (also named mercaptans), sulfur compounds (hydrogen or ammonium sulfide), low-molecular-weight carboxylic acid also known as short chain fatty acids (as propionic or butyric acid), aldehydes, amines and heterocyclic compounds. Odorant, according to their chemical class, combination and/or concentration, lead to good or unpleasant odors^1,2^. Among the several gut microbial odorants and volatile organic compounds, butyrate is of particular interest to human health as it is an energy source for colonic cells. In addition, it inhibits inflammation and carcinogenesis, reinforces colonic defense barrier, modulates gut epithelium permeability, promotes satiety, increases sodium and water reabsorption (anti-diarrheal agent) and reduces oxidative stress^3^. Production of butyrate by gut commensals drive the immunity of the host by inducing the differentiation of colonic and systemic regulatory T cells,^4,5^ and has an important epigenetic effect on the gut epithelial cells by inhibiting histone deacetylase (HDAC)^6,7^. More specifically, a lower butyrate to acetate ratio has been associated with non-alcoholic steatohepatitis (NASH), adenomatous polyps or colon cancer^8^.

On the other hand, we have recently identified a link between redox state and microbiome^9^. That is, a link between redox potential, oxidative stress and human microbiota according to the oxygen tolerance of each species and the abundance of antioxidants in the environment. For the first time, we found a statistically significant association between fecal redox potential and metagenomic relative aerotolerant predominance^9^. In severe malnutrition, the lack of antioxidants and nutrients necessary for endogenous antioxidants and a major liver peroxisomal and mitochondrial dysfunction results in a major oxidative stress^10–12^. In children with severe acute malnutrition, we have shown a major depletion of anaerobic microbes (or intolerant to reactive oxidative species)^12^, consistent with the collapse of fecal butyrate in those who die^13^. This led us to found that antioxidants allow most anaerobes, including methanogenic archaea, to thrive in an oxidative environment (aerobic)^14–16^.

Serendipitously, we observed an absence of unpleasant odor when anaerobes were grown aerobically with antioxidants. To test what might explain a better smell, we selected 3 strains of the human gut isolated by microbial culturomics^17–20^, that produced the strongest smells in our anaerobic culture laboratory: *Clostridium sporogenes, Clostridium subterminale* and *Romboutsia lituseburensis*. We compared the volatile organic compounds (VOCs) of these 3 strains grown under standard anaerobic condition or in ambient air thanks to antioxidants (uric acid, ascorbate, glutathione)^14–16^ using gas chromatography coupled with mass spectrometry. We were surprised to see that the profile of volatile organic compounds was profoundly altered by the atmosphere. Aerobic metabolic repertoire was different from the anaerobic repertoire and this has been confirmed by analysis of polar metabolites. Anaerobic but not aerobic repertoire allowed to discriminate each species with perfect accuracy. Surprisingly and unexpectedly, butyrate production was increased aerobically with antioxidants, suggesting a critical role of antioxidants in the gut mucosa for the maintenance of butyrate production.

## MATERIAL AND METHODS

### Aim, Design and Setting of the Study

The aim of the study was to explore the metabolism switching of human gut anaerobes isolated by microbial culturomics^17–20^ producing volatiles organic compounds when grown in anaerobiosis or in aerobiosis with antioxidants. The design was an *in vitro* experimental study on human gut strains. The setting was our center, specialized in the human microbiome and where the approach of microbial culturomics was set up since 2012 with analysis of more than 1,000 human samples allowing us to isolate more than 1,600 bacterial species from human gut and more than 360 new human species to date^17–20^ allowing us to substantially extend the human gut repertoire^20^.

### Description of materials

#### Culture conditions

Three strains of anaerobic bacteria, isolated by culturomics studies^17–20^ and obtained from the Collection de Souches de l’Unité des Rickettsies (CSUR, Marseille, France), were used for this study: *Clostridium sporogenes* CSUR P3957, *Clostridium lituseburense* CSUR P3909 and *Clostridium subterminale* CSUR P3759. Those bacterial strains were previously cultured 24 hours anaerobically at 37 °C on Columbia agar supplemented with 5% of sheep blood (bioMérieux, Marcy l’Etoile, France). Then, the bacterial inoculum was calibrated to 10^6^ bacteria/mL in PBS using a densitometer (Thermo Ficher, Villebon sur Yvette, France) and 0.1 mL of this solution (10^5^) was inoculated into the different culture bottles using a syringe. Classical anaerobic culture condition in standard anaerobic blood culture (Becton Dickinson, Le Pont de Claix, France) under 100% nitrogen atmosphere was compared with the aerobic culture supplemented by the antioxidant cocktail in aerobic blood culture bottles (Becton Dickinson). Negative culture controls were performed under classical aerobic condition in aerobic blood culture bottles (Becton Dickinson) without antioxidants. 5% sheep blood was previously added in each culture conditions. The antioxidant solution was prepared as follow: a stock solution containing 4% ascorbic acid (VWR, Leuven, Belgium), 0.4% glutathione and 2% Uric Acid (Sigma-Aldrich, Lyon, France) was prepared in distilled water and filtered using a 0.2 μm filter after adjusting pH to 7.5 with KOH 10M. Then, 1 mL of this solution was injected into the aerobic blood culture bottle (Becton Dickinson, Le Pont de Claix, France) containing 40 mL of culture medium supplied by the manufacturer (for a final concentration of antioxidant of 1 g/L ascorbic acid, 0.1 g/L glutathione and 0.4 g/L uric acid) using a sterile syringe and needle to avoid contamination. All the blood culture bottles were incubated at 37°C for different time (48 to 96h, see behind) before measure of VOC, SCFA and polar metabolites.

To confirm the viability of the bacterial strains at the end of the experiment, the blood culture content was subcultured anaerobically for 48 hours on Columbia Agar medium supplemented with 5% sheep blood (bioMérieux). Then, the identification of the three bacterial strains was confirmed by MALDI-TOF-MS as previously described^21^. No viable bacteria have been isolated after aerobic culture, while viability and growth of all strains were confirmed in aerobic blood culture bottles supplemented with antioxidant and in anaerobic blood culture bottles.

#### Volatile Organic Compounds (VOCs) investigation

To measure the temporal production of VOCs in the bottles, different times of incubation were tested for each culture condition: 48, 72 and 96 hours at 37 °C. After selecting the most suitable incubation time of 72 hours, VOCs were measured in triplicate for each condition. VOCs were collected for 10 minutes using a PDMS/DVB 65 μm SPME fiber (Supelco, Sigma-Aldrich, Saint-Quentin Fallavier, France) inserted in the headspace of each culture bottle. Gaz chromatography-Mass spectrometry (GC/MS) analyses were carried out on a Clarus 500 gas chromatograph equipped with a SQ8S MS detector (Perkin Elmer, Courtaboeuf, France). VOCs were volatized during 15 minutes from the SPME fiber at 250°C (splitless, 1 mm internal diameter liner) and separated on an Elite-5MS column (30 m, 0.25 mm i.d., 0.25 mm film thickness) using a temperature gradient: 50 °C (5 min) to 100 °C at 4°C/min, 100 to 250 °C at 30 °C/min. Helium flowing at 1 mL/min was used as carrier gas. MS inlet line was set at 250 °C and Electron Ionization source at 200 °C. Full scan monitoring was performed from 33 to 500 m/z. All data was collected and processed using Turbomass 6.1 (Perkin Elmer, Courtaboeuf, France). Spectral database search was performed using MS Search 2.0 operated with the Standard Reference Database 1A (NIST, Gaithersburg, USA). Chemical identifications were validated with both Reverse and Forward scores above 800. Retention indexes were used to confirm the identifications. The area under the peak curve was then collected for each identified peak. The measurement of a blank sample was carried out in parallel, consisting of a non-inoculated blood culture bottle containing the same culture medium as the inoculated bottles. Consequently, results only consider VOCs measured in culture samples after subtracting the corresponding blank values.

#### Non-Esterified Short chain fatty acids (SCFAs) quantification

After finding that non-esterified short-chain fatty acids (SCFA) were not detected by our first approach, we measured them with an appropriate specific technique that detected acetic acid (C2) to heptanoic acid (C7). Acetic, propanoic, isobutanoic (isobutyric), butanoic (butyric), isopentanoic (isovaleric), pentanoic (valeric), hexanoic (caproic) and heptanoic acids were purchased from Sigma Aldrich (Lyon, France). A stock solution was prepared in water/methanol (50% v/v) at a final concentration of 50 mmol/L and then stored at –20 °C. Calibration standards were freshly prepared in acidified water (pH 2-3 with HCl 37%) from the stock solution at the following concentrations: 0.5, 1, 5 and 10 mmol/L. SCFAs were analyzed from 3 independent culture bottles (both blank and samples). Culture medium was collected and then centrifuged 5 minutes at 16,000 x g to remove bacteria and debris. The clear supernatant was adjusted to pH 2-3 and spiked with 2-ethylbutyric acid as the internal standard (IS) at a final concentration of 1 mmol/L (Sigma Aldrich, Lyon, France). The solution was centrifuged again before injection. SCFAs were measured with a Clarus 500 chromatography system connected to a SQ8s mass spectrometer (Perkin Elmer, Courtaboeuf, France) such as detailed previously^22^. Aqueous samples were directly injected (0.5 µL) in a splitless liner heated at 200 °C. Injection carry-over was decreased with 10 syringe washes in methanol: water (50:50 v/v). Compounds were then separated on an Elite-FFAP column (30 m, 0.25 mm i.d., 0.25 mm film thickness) using a linear temperature gradient from 100 to 200 °C at 8 °C/min. Helium flowing at 1 mL/min was used as carrier gas. MS inlet line and Electron Ionization source were set at 200 °C. To insure compound selectivity, Selected Ion Recording (SIR) was performed after a 4.5 min solvent delay with the following masses: 43 m/z (isobutanoic acid), 60 m/z (acetic, butanoic, pentanoic, isopentanoic, hexanoic and heptanoic acids) 74 m/z (propanoic acid), 88 m/z (2-ethylbutyric acid, IS). All data was collected and processed using Turbomass 6.1 (Perkin Elmer, Courtaboeuf, France). Quadratic internal calibration was calculated for each acid using the peak areas from the associated SIR chromatograms. Coefficients of determination were all above 0.999. Back-calculated standards and calculated quality controls (0.5 and 5 mmol/L) all showed good accuracy with deviations below 15%. SCFAs quantities in culture samples were presented after subtraction of the quantities measured in the blank samples (average production of 2 non-inoculated aerobic blood cultures). The upper limit concentration detectable by the system was 10mM.

### Polar metabolites investigation

#### Sample preparation

*Clostridium sporogenes* was cultivated under the same conditions as described above. Five culture replicates for the anaerobic and the aerobic antioxidants cocktail culture conditions were prepared, as well as their five respective culture controls (without bacteria). 1 mL of each culture sample was centrifuged (6,000 × g, 10 minutes, 4 °C). The collected cell pellets were re-suspended in 400 mL of methanol at −80°C to quench the sample and tubes were vortexed 2 minutes. Solutions were incubated during 30 min at −80°C and then centrifuged again (14,000 × g, 10 minutes, 4 °C). The supernatants were dried under a stream of nitrogen at room temperature (Liebish, Thermo Fisher Scientific, Courtaboeuf, France) and stored at - 80°C until LC/MS analysis. Dry samples were reconstituted in 50 µL of 10% acetonitrile, 90% water and 0.1% formic acid (ULC/MS grade solvents; Biosolve Chimie, Dieuze, France). Pooled Quality Control (QC) samples were prepared by pooling all bacteria samples with equivalent quantities (culture controls were not pooled).

#### LC/MS analysis

Metabolites were first separated by reverse phase chromatography using an ACQUITY UPLC® I-Class System (Waters, Saint-Quentin-en-Yvelines, France) and then monitored using a hybrid Ion Mobility QTOF mass spectrometer (Vion, Waters, Saint-Quentin-en-Yvelines, France). Samples were stored at 10 °C during analysis and injected following a random sample list including pooled QC samples every 5 injected samples. 5 µL of reconstituted samples were separated on a reverse phase HSS T3 column (1.8 µm, 2.1 × 100 mm, Waters, Saint-Quentin-en-Yvelines, France) thermostated at 50 °C. Metabolites were eluted through the column at 0.5 mL/min using the following solvent composition gradient: 1% acetonitrile 0.1 % formic acid in water 0.1 % formic acid during 0.5 minutes, 1 to 20% during 5.5 minutes, 20 to 80% during 4 minutes followed by a 1.5 minutes wash step a 98% and then a 3.5 minutes equilibration step towards initial conditions. Total run time was 15 min. Compounds were ionized in a Zspray ESI source using a default “Labile Tune” to reduce compound fragmentation before QTOF monitoring. Ionization parameters were set as follow: 0.8/2.0 kV for positive/negative modes, source/desolvation temperatures = 120/600 °C, cone/desolvation gaz = 100/600 L/h. Ions were monitored using a Sensitivity HDMSe data independent analysis method between 50 and 1000 m/z as follow: scan time 0.15 s, collision energy ramp 20-40 eV, automatic mass correction during survey (using Lockmass 554.2620 m/z from a Leucine Enkephalin solution). All ions produced in-source were alternatively analyzed at low/high fragmentation energy to collect parent/fragments masses throughout the analysis. Mass and CCS calibrations were run before analysis batches using the Major Mix solution (Waters, Saint-Quentin-en-Yvelines, France). Instrument performance were checked before, during and after analysis batches using a standard mixture (System Suitability Test Quality Control, Waters, Saint-Quentin-en-Yvelines, France). Mass errors and CCS deviations were below 5 ppm and 2% when injected alone or mixed with the sample pooled QC. No significant performance changes were noticed during the analysis batches.

#### Data processing

Raw data was then processed by UNIFI software to collect “components” described by a retention time, a drift time and parent/fragment masses. Automatic parameters were set to detect 4D peaks above 150 and 75 counts for low and high energy scans. Once all components were listed, a marker table was generated with the following parameters: 0.2-12 minutes retention time window, intensity above 2000 counts, automatic mass/ drift time/ retention time tolerances (14.2 ppm/ 0.47 ms / 0.03 min for positive experiment and 16.2 ppm/ 0.54 ms/ 0.03 min for negative experiment). The marker table was sent to the EZinfo multivariate statistics software (Umetrics, Umea, Sweden) for data exploration by PCA then data classification according to both anaerobic and aerobic groups using an OPLS-DA model. From the OPLS-DA model, the VIP and S-plot diagrams were used to select discriminant markers as follow: VIP>1 and S-plot markers >90%. The selected markers were then labelled on the marker table in UNIFI and searched using the elucidate tool against the following structure libraries: metabolite structures (including CCS values, Waters, Saint-Quentin-en-Yvelines, France), then KEGG+HMDB databases (Chemspider access). Potential identities were then added to an inhouse structure library including the structure .mol files from Chemspider (Royal Society of Chemistry, London, United Kingdom) and then targeted on the initial raw data according to the following parameters: 5 ppm mass tolerance, 2 mDa mass tolerance for structure predicted fragments, 5% CCS deviation, multiple adduct finder (+H, +Na, +K, +Li, +NH4) within 5 ppm and 0.1 minutes. Identifications were then filtered with the following parameters in order to validate metabolites: mass error < 2 ppm on parent ions, < 2% CCS deviation (if available), at least one predicted structure fragment, < 10% error on CCS estimates, < 5 ppm error on RMS isotope match and < 15% RMS percent isotope intensity.^23^ Each component was validated across all injections according to: equivalent retention time to that form the marker table, isomers with equivalent retention times and CCS values, summary plot showing different responses between both culture conditions. Stringent filters were removed to collect the response of valid metabolites across all samples. Metabolites were also checked to be consistent between positive and negative experiments (retention time, CCS values, fold changes). The fold change was defined as the ratio of response values from the mean response for each condition.

### Statistical analysis

The peak areas of each detected VOC (arbitrary unit) and the concentration of each SCFA (mM) were assessed as quantitative variables. Species (*C. sporogenes, R. lituseburensis* and *C. subterminale*) and atmosphere (aerobiosis without antioxidant, aerobiosis with antioxidant and anaerobiosis) were considered as qualitative variables. Except mentioned otherwise, three replicate values were measured. Comparing aerobiosis with antioxidant and anaerobiosis, three t-tests (one by species) were performed, and discoveries were determined using the two-stage linear step-up procedure of Benjamini, Krieger and Yekutieli, with Q = 1% (VOC) or 5% (SCFA). Each row was individually analyzed, without assuming a consistent SD. When 3 groups were compared (aerobiosis without antioxidants, aerobiosis with antioxidants and anaerobiosis), two-way ANOVA was also performed with correction for multiple comparisons using the Tukey’s test. Principal component analysis with the Pearson metrics was performed and biplot were shown to explore the association between strains, atmosphere and metabolites. Statistical tests were two-sided. A p-value ≤ 0.05 was considered significant. These statistical analyses were performed using GraphPad v8.0 (GraphPad Software, La Jolla, CA, USA) and XLSTAT v19.1 software (Addinsoft, Paris, France).

## RESULTS

### Amount and diversity of volatile organic compounds

The number and total amount of VOCs was higher in anaerobiosis for the 3 species and the difference was more significant considering only odorous VOCs (Fig. 1). Both atmosphere and species determined the diversity and quantity of VOCs and odorous VOCs. However, the magnitude of the atmosphere effect was more important on VOCs amount than on their diversity (number of VOCs increased 1.2- to 1.8-fold (odorous VOCs 2.0- to 3.7-fold), while the total amount was increased 5.1- to 7.4-fold (7.7- to 13.6-fold) in anaerobiosis). The maximal atmosphere effect was observed for the total amount of odorous VOCs for *Clostridium sporogenes* (13.6-fold ratio, p-value < .00005, q-value < .0005).

**Figure 1.**
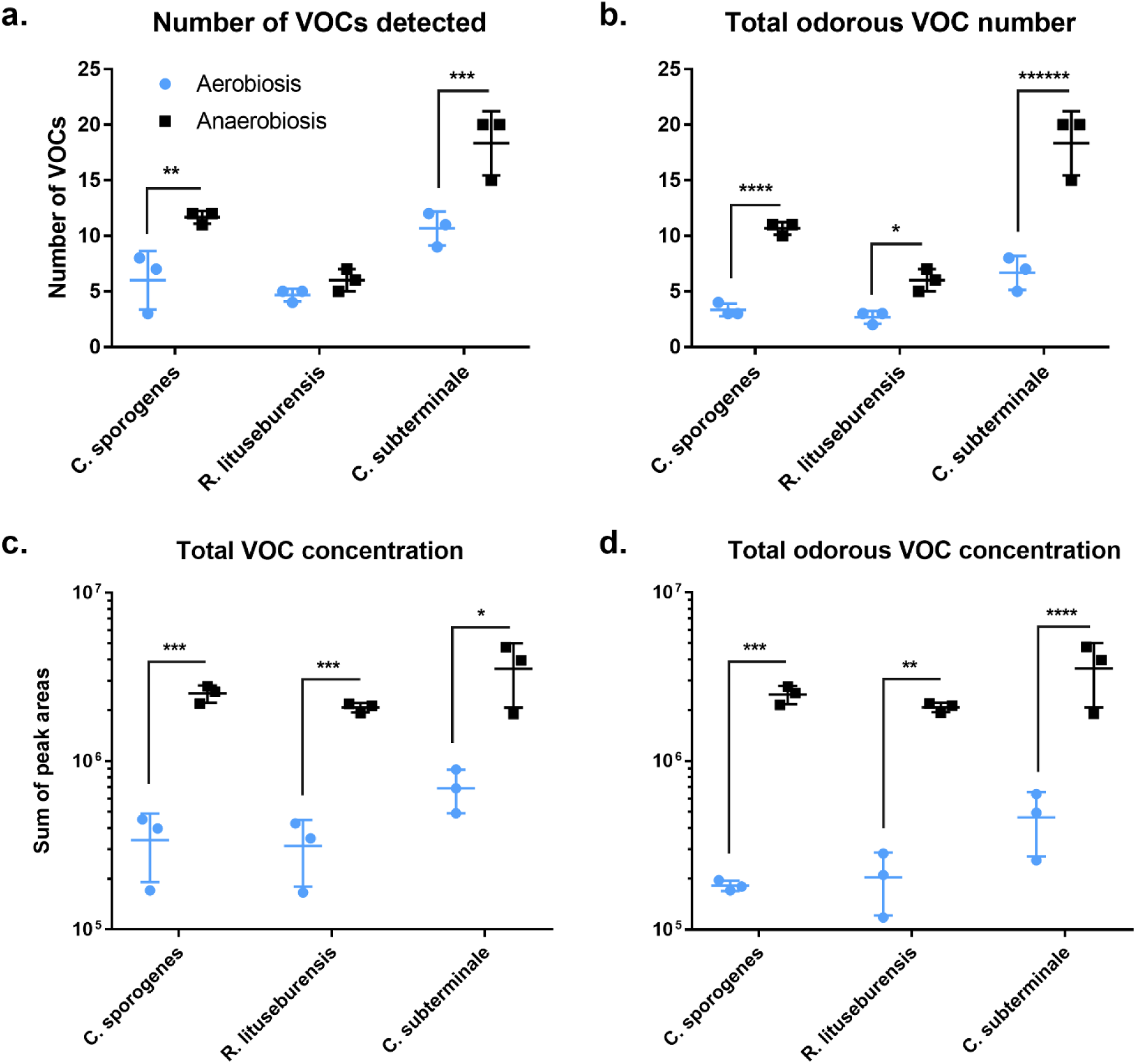
Diversity and total concentration of volatile organic compounds under antioxidant-based aerobic or anaerobic culture condition. VOCs: volatile organic compounds. The number (**a**) and total amount (**c**) of VOCs was higher in anaerobiosis for the 3 species and the difference was more significant considering only odorous VOCs (**b**,**d**). *p<0.05, **p<0.005, and so on.

### Chemical classes of volatile organic compounds

The 37 different VOCs produced by the 3 strains corresponded to 6 chemical classes, including 15 short chain fatty-acid ester (SCFAE), 7 sulfur compounds (thiols), 7 alcohols, 11 alkanes, 2 aromatic compounds and 1 alkene (Fig. 2, Table S1). By analyzing the number of VOCs detected for each culture condition regardless of the species, we found that the diversity of VOCs was reduced in aerobiosis since 17 VOCs were detected in aerobiosis compared to 28 in anaerobiosis (p <.05). Alkanes, aromatic compounds, alcohols, short chain fatty acid esters, sulfur compounds were detected both in aerobiosis and anaerobiosis (Fig. 2b). The alkene (1,3,5-cycloheptatriene) was detected only in aerobiosis. Sulfur alcohols and sulfur SCFAE were detected only in anaerobiosis. Only one sulfur compound ((methyldisulfanyl) methane, commonly named dimethyl disulfide) and one SCFAE (3-methylbutyl 3-methybutanoate) were detected both in aero and anaerobiosis. This pattern remains unchanged over time as observed by analyzing VOCs produced at 48, 72 and 96 hours (Figure S1).

**Figure 2.**
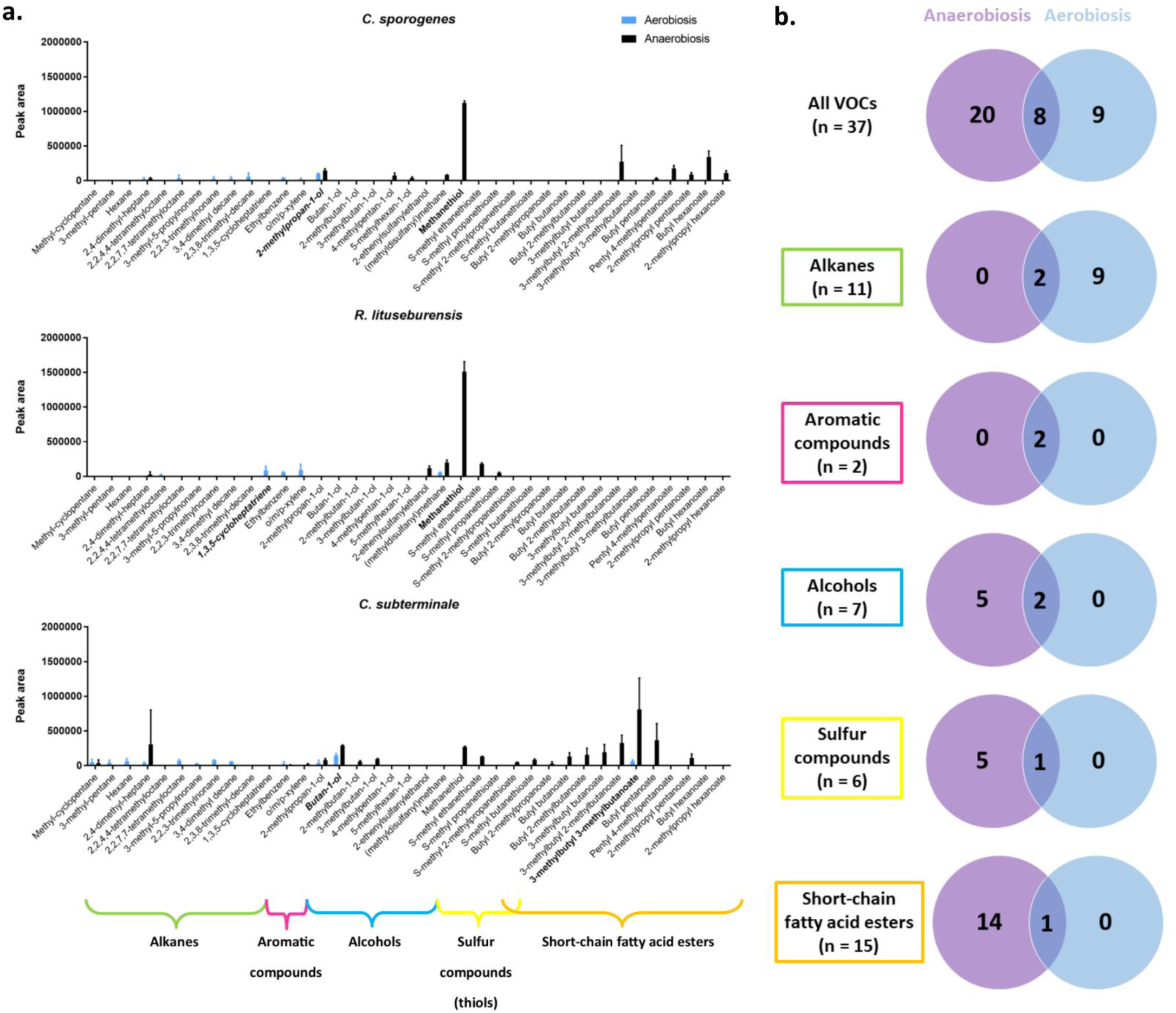
Diversity of volatile organic compounds according to species, chemical class and antioxidant-based aerobic or anaerobic culture condition. VOCs: volatile organic compounds. The 37 different VOCs produced by the 3 strains corresponded to 6 chemical classes, including 15 short chain fatty-acid ester (SCFAE), 7 sulfur compounds (thiols), 7 alcohols, 11 alkanes, 2 aromatic compounds and 1 alkene (**a**,**b**). 9 VOCs (alkanes) were detected only in aerobiosis (**b**). Anaerobiosis was associated with a specific and exceptional production of methanethiol for *R. lituseburensis* and *C. sporogenes* (**a**). The aerobic metabolic repertoire is different from the anaerobic one and is restricted suggesting a constrained metabolic behavior for VOC production redirected to the alkanes in the presence of oxygen.

This unexpected metabolic switching was confirmed by principal component analysis (Fig. 3). Indeed, the VOCs associated with aerobic culture with antioxidants were clearly different and clustered compared to the VOCs produced anaerobically. In aerobiosis with antioxidants, the 3 bacterial strains were found adjacent and the VOCs produced mainly belong to 2 classes, including 10 alkanes and 2 aromatic compounds (o/m/p-xylene and ethylbenzene). In contrast, VOC produced anaerobically exhibited a greater heterogeneity. *C. sporogenes* and *R. lituseburensis* were grouped together, whereas *C. subterminale* was clearly different. This contrasts with the phylogenetic analysis (Figure S2). However, despite this heterogeneity, only one alkane was associated with anaerobic culture (2,4-dimethyl-heptane by *Clostridium subterminale*), while all short-chain fatty acid esters, thiols and alcohols were associated with anaerobiosis (Fig. 2 & 3). This suggests that there is not only a quantitative change between the atmosphere but that a specific and unexpected aerobic metabolic repertoire can be observed when the 3 *Clostridia* species are grown in aerobiosis with antioxidants. This aerobic metabolic repertoire is globally different from the anaerobic one.

**Figure 3.**
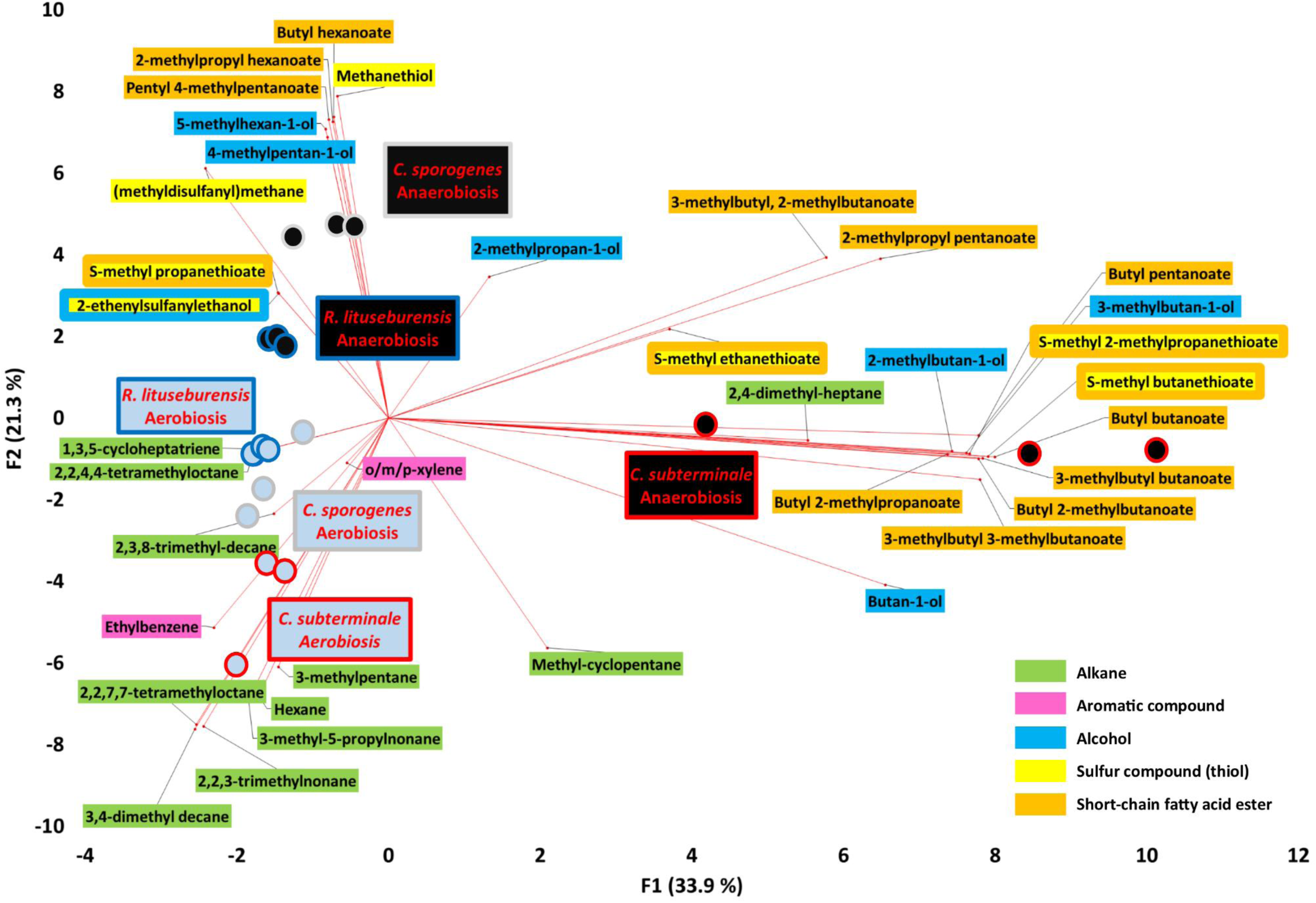
Biplot of the principal component analysis showing associations between species, culture conditions and volatile organic compounds. The VOCs associated with aerobic culture with antioxidants were clearly different and clustered compared to the VOCs produced anaerobically. In aerobiosis with antioxidants, the 3 bacterial strains were found adjacent and the VOCs produced belong to 2 classes, including 10 alkanes and 2 aromatic compounds (o/m/p-xylene and ethylbenzene). In contrast, VOC produced anaerobically exhibited a greater heterogeneity. This suggests that there is not only a quantitative change between the atmosphere but that a specific and unexpected aerobic metabolic repertoire can be observed when the 3 *Clostridia* species are grown in aerobiosis with antioxidants. Principal component analysis performed using XLSTAT v19.1.

### Restricted metabolic repertoire in aerobiosis

By analyzing the amount of each VOC per species (Fig. 2 & Fig. S3, Table S1), we observed that anaerobiosis was associated with a specific and exceptional production of methanethiol for *R. lituseburensis* (7.5-fold the second VOC - (methyldisulfanyl)methane, commonly called dimethyl disulfide) and *C. sporogenes* (3.2-fold the second VOC production - butyl heaxanoate). For *C. subterminale*, 3-methylbutyl 3-methylbutanoate was the most anaerobically produced VOC (Fig. 2 & Fig. S3). We observed that *R. lituseburensis* produced neither SCFAE nor alcohol and that *C. sporogenes* produced no sulfur SCFAE whatever the atmosphere. In contrast, *C. subterminale* produced members of all VOC classes, including several SCFAE but little anaerobic methanethiol compared to the other two species. This suggests that the anaerobic VOC repertoire of *C. subterminale* is more diverse than that of *C. sporogenes* and *R. lituseburensis*. In comparison, the aerobic metabolic repertoire is restricted and more similar among the 3 species, suggesting a constrained metabolic behavior for VOC production in presence of oxygen. A heatmap unsupervised analysis (Fig. 4) confirmed the dichotomic metabolic switching since the atmosphere was the main discriminant factor and had a larger effect that species. Moreover, the accuracy of species identification was better in anaerobiosis (perfect accuracy by heatmap unsupervised analysis, Fig. 4) and this was confirmed by discriminant analysis with 87.5% correct results in anaerobiosis versus 75% in aerobiosis after cross-validation.

**Figure 4.**
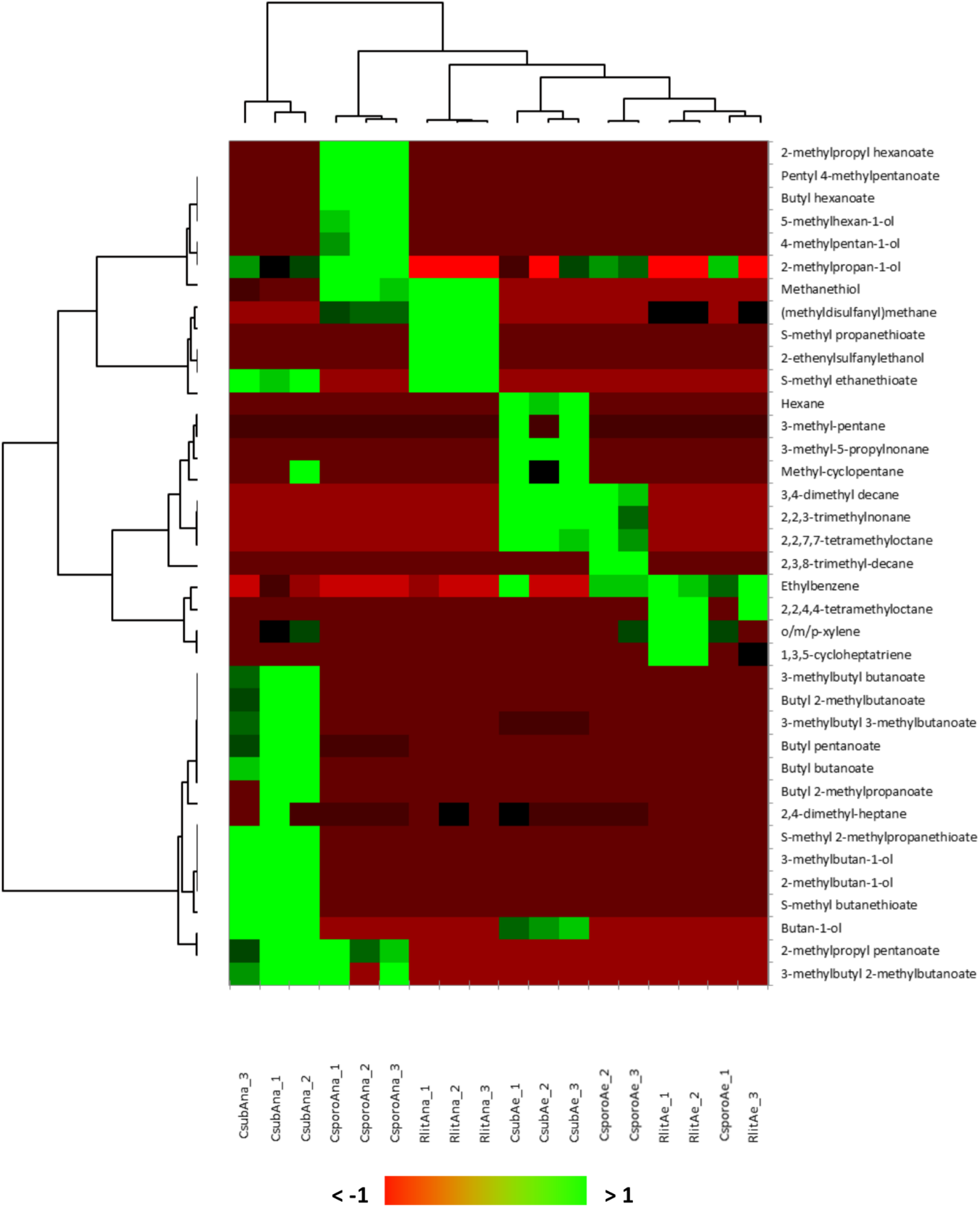
Species-specific VOC profile in anaerobiosis than in aerobiosis. A heatmap unsupervised analysis confirmed the dichotomic metabolic switching based on VOCs production since the atmosphere was the main discriminant factor and had a larger effect that species. Moreover, the accuracy of species identification was better in anaerobiosis (branch length between species was longer in anaerobiosis than in aerobiosis with an aerobic classification error: *C. sporogenes* among *R. lituseburensis –* perfect classification in anaerobiosis). The better species resolution in anaerobiosis was confirmed by discriminant analysis with 87.5% correct results in anaerobiosis versus 75% in aerobiosis after cross-validation. Heatmap unsupervised analysis performed using XLSTAT v19.1. Each experiment was replicated 3 times.

### Butyrate production

We subsequently tested the production on non-esterified short chain fatty acids in the presence of oxygen as near the gut epithelium. Beyond the critical role of antioxidant to preserve the gut epithelial associated “anaerobic” microbes^16,24^, this could be a proof of concept for the critical role of gut antioxidant capacities to maintain the production of beneficial molecules by “anaerobes”, as butyrate under hyperoxygenation.^25,26^ Very minimal acetic acid production was detected in vials of aerobic blood cultures without supplementation with antioxidants, while all other SCFA appeared to be consumed even though the difference in concentration was very small (Fig. 5). Since none of the 3 bacterial strains survived in aerobiosis in the absence of antioxidants, this could represent a transient metabolism before bacterial death. Pentanoic, hexanoic and heptanoic acid were not produced by any strain in any atmosphere. Acetic acid (C2) was very abundantly (>10mM: upper detection limit) produced by all strains in aerobic blood culture bottles supplemented with antioxidants, and in anaerobiosis. A significant difference according to both atmosphere and species was confirmed by two-way ANOVA (p < .0001) for propanoic, butanoic, isobutanoic and isopentanoic acids with a very significant interaction. By comparing only aerobiosis with antioxidants to anaerobiosis by multiple t-tests (one for each species) corrected for multiple comparisons, we were surprised to find that SCFA production was often higher in aerobiosis with antioxidants than in anaerobiosis (Table S2). This was particularly significant for *R. lituseburensis. C. sporogenes* which produced more isobutanoic than butanoic acid. *C. subterminale* did not produce a significant amount of propanoic acid, but was the best producer of butanoic and isopentanoic acid. Indeed, it produced up to 8.7 ± 1.0 mM of butanoic acid and 7.5 ± 1.6 mM of isopentanoic acid in aerobiosis with antioxidants (Fig. 5 & Table S2).

**Figure 5.**
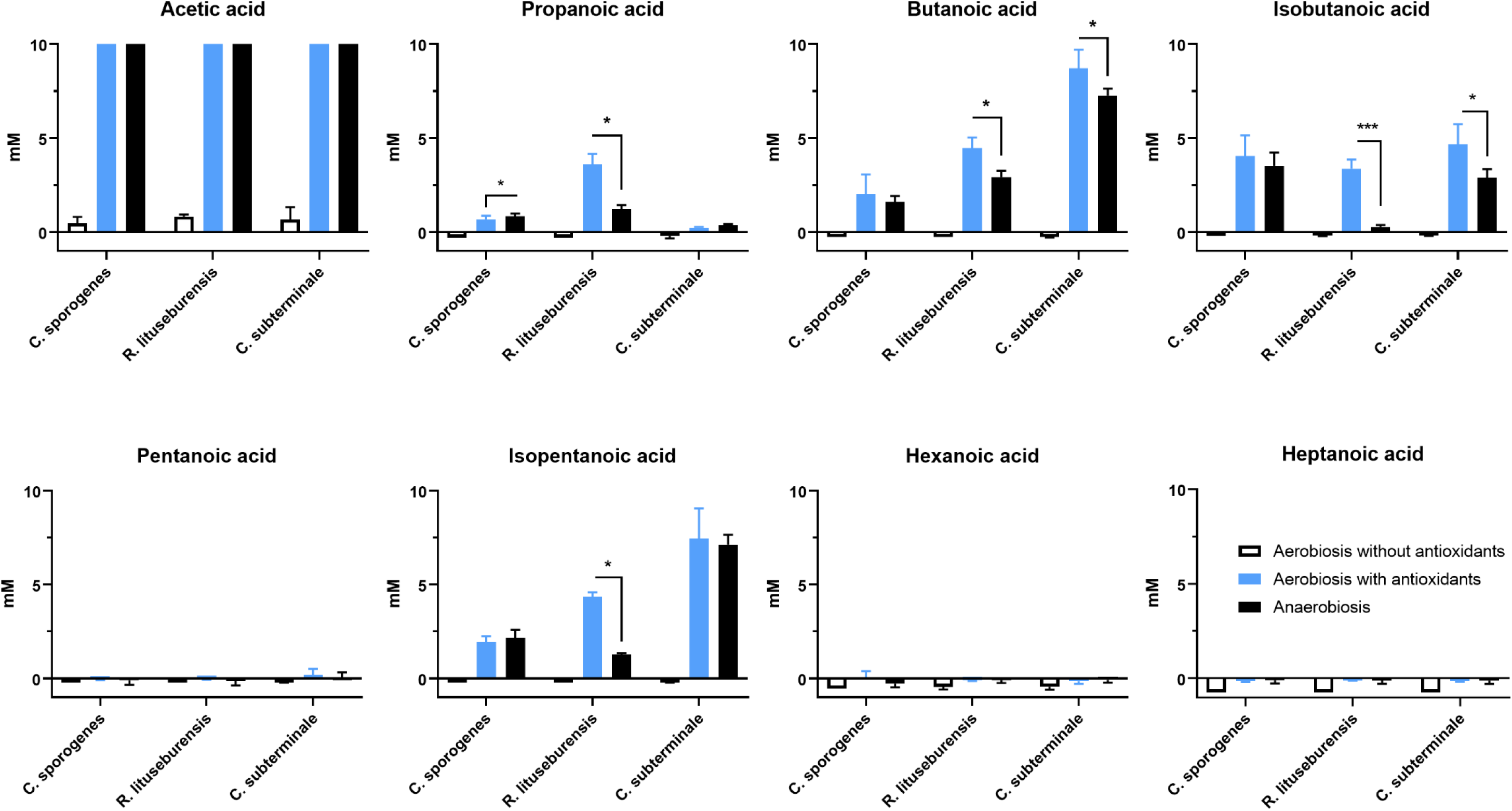
Comparison of short-chain fatty acids produced in antioxidant-based aerobic or anaerobic culture condition. Acetic acid (C2) was very abundantly (>10mM: upper detection limit) produced by all strains in aerobic blood culture bottles supplemented with antioxidants, and in anaerobiosis. Production butanoic acid (butyrate), a key metabolite for human gut homeostasis, was higher in aerobiosis with antioxidants than in anaerobiosis. *p<0.05, ***p<.0005.

### Polar metabolites

By analyzing the production of polar metabolites (Fig. 6), we observed that there was a different profile in aerobiosis, with a major increase in the production of many lysophospholipids and some dipeptides. The detection of oxidized glutathione in aerobic conditions confirmed its antioxidant activity in the culture medium. On the other hand, pilocarpine, a parasympathicomimetic specific for M3 muscarinic receptors that promote digestive peristalsis, was increased anaerobically. Some metabolites were specific to each atmosphere: 3’-uridylic acid, 4-hydroxy-1-pyrroline-2-carboxyclic acid and lysophophatidylethanolamine (18:1n7/0:0) were detected only in aerobic conditions (Table S3). 1,4’-bipiperidine-1’-carboxyclic acid, 3-hydroxy-8’-apo-E-caroten-8’-al, Cyclo(L-Trp L-Pro), heptanal, 2-benzylidene-(E), leucine proline dipeptide, meconin, and SHCHC (2-succinyl-6-hydroxy-2,4-cyclohexadiene-1-carboxylate, precursor of vitamin K) were detected only anaerobically (Table S4).

**Figure 6.**
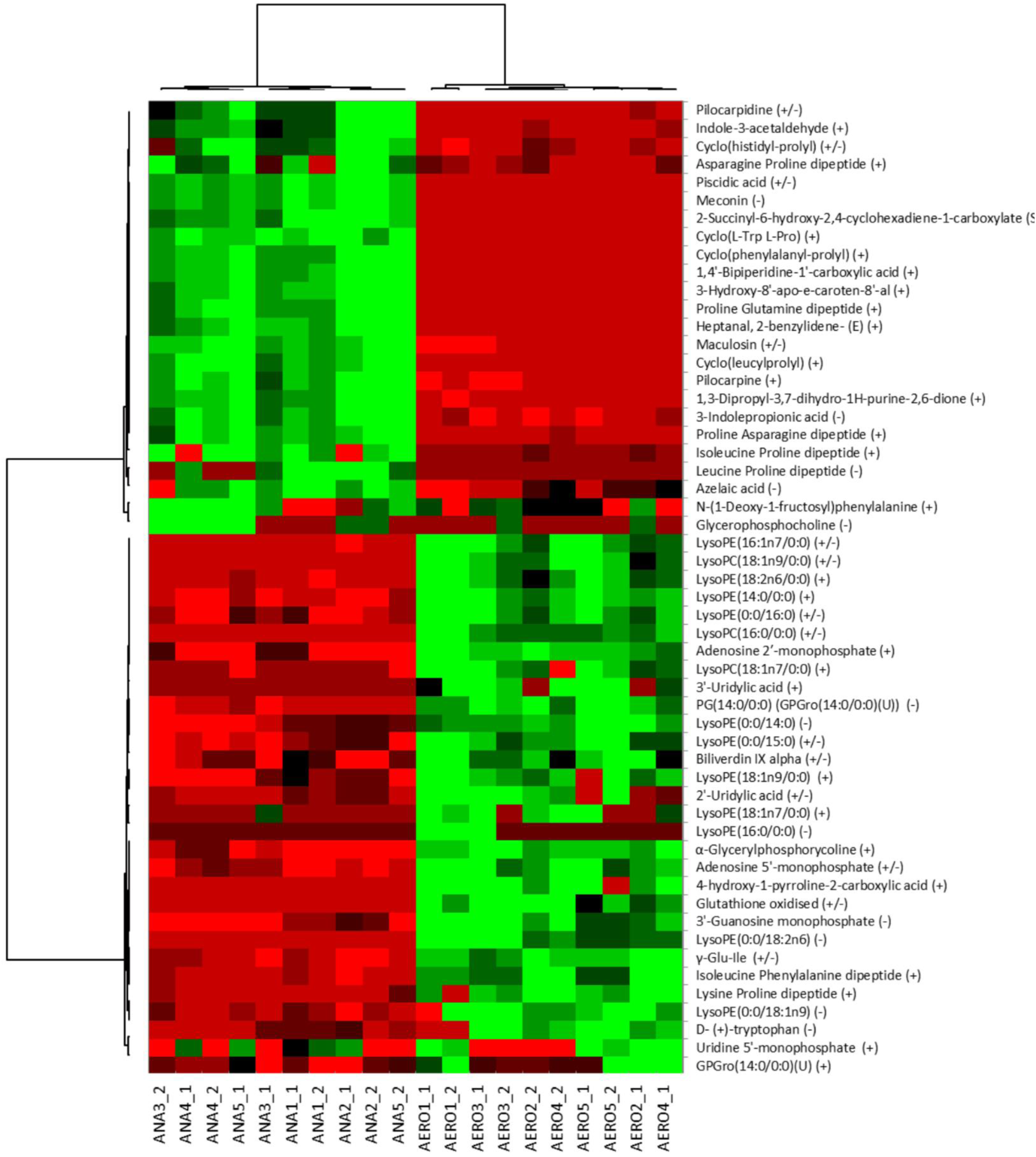
Polar metabolite profile according to atmosphere. The profile of polar metabolites was different in aerobiosis, with a major increase in the production of many lysophospholipids and some dipeptides. The detection of oxidized glutathione in aerobic conditions confirmed its antioxidant activity in the culture medium. ANA: Anaerobiosis, AE: Aerobiosis. This analysis was performed only for *C. sporogenes*. Measurements were performed twice for each of 5 experiments. Heatmap unsupervised analysis performed with XLSTAT v19.1.

## DISCUSSION

Here, we have shown the metabolomic switching when human gut butyrate-producing *Clostridia* are exposed to oxygen in the presence of antioxidants and the critical role of antioxidants in maintaining butyrate production in this context. The oxygen level is a critical biochemical parameter determining the composition of gut microbiota^27,28^. In the distal gut lumen, anaerobiosis is obtained by the consumption of oxygen, either by digestive microbes, by the epithelium or even by non-microbial reactions such as lipid oxidation^28^. Luminal oxygen levels in the lumen of the distal small intestine (terminal ileum) are near zero (10 mmHg) and much lower than in the proximal digestive tract (60mmHg)^28^. Enteric pathogens, such as *Citrobacter rodentium* or *Salmonella enterica* are able to modify the gut microenvironment themselves by increasing the oxygenation of the mucosal surface via mucosal hyperplasia caused by their type III secretion system (for *C. rodentium*^26^) or by depletion of the *Clostridia* that produce butyrate (for *S. enterica*^25^) to promote their aerobic growth. Indeed, aerobic respiration causes uncontrolled luminal expansion of the pathogen^26^. Consequently, digestive hyperoxygenation is a pathogenicity mechanism of enteric pathogens that can disrupt host-archaeal-bacterial mutualism.

Anaerobiosis rapidly alters the metabolism of facultative anaerobes,^29^ However, this has never been studied for strict anaerobic bacteria because until recently, it was considered that their growth in the presence of oxygen was not possible^14–16^. We observed that the metabolism of the 3 species studied was restricted and redirected to different metabolic pathways in aerobiosis versus anaerobiosis, thus explaining the absence of typical and strong smell in aerobiosis. The conservation of butyrate production in aerobiosis with antioxidants observed here is particularly important because butyrate itself limits the oxygenation of digestive lumen; in the colonocytes, oxygen consumption with oxidation of butyrate by colonocytes gives CO2 and ATP, and this is associated with water and sodium absorption^25^. Our findings shed light on the beneficial role of antioxidants on gut microbiota and butyrate production, previously reported in vivo, particularly with oligomeric proanthocyanidins (OPC)^30,31^ but microbial mechanism was not understood. Here, we show that antioxidants unravel an unexpected and unexplored aerobic metabolic repertoire of the human gut *Clostridia*, and this repertoire is possibly active in vivo near the human gut mucosa. The physiological effect of the possible microbial production of alkanes, alkenes, and lysophospholipids near the human gut mucosa will need to be clarified in future studies. Nevertheless, the effect of butyrate on human gut mucosa and immunity is well known and is key to human health^4,5,32^.

## CONCLUSIONS

Here, we demonstrated that a combination of the 3 major human antioxidants (ascorbic acid, glutathione and uric acid) allow human gut *Clostridia* to thrive and maintain microbial production of the key molecule butyrate in the presence of oxygen (in aerobiosis). This work provides new arguments for the understanding of the critical role of antioxidants in the maintenance of the gut homeostasis and host-archaeal-bacteria mutualism and its resilience to oxidative aggression.

## DECLARATIONS

### Additional files

Additional file 1. Supplementary figures and tables. This file contains supplementary Figures S1-S4 and Tables S1-S4.

#### Acknowledgements

We would like to thank all the culturomics team for providing human gut strains.

### Ethics approval and consent to participate

Not applicable

### Consent for publication

Not applicable

### Availability of data and material

The data that support the findings of this study are available from the corresponding author.

### Authors’ contributions

M.M. and D.R. conceived the main idea, and S.K. and N.A. proved it. S.K., G.D. and E.G. performed the microbial culture. N.A., E.G., M.R. and E.C. performed the metabolic analyses. M.M., G.D, J-C.L and D.R. extensively analyzed and interpreted the results and contributed to the writing of the manuscript. E.C. and D.R. designed the study and supervised the work.

