## Supplementary data for "Antioxidants maintain Butyrate production by Human Gut *Clostridia* in the presence of Oxygen"

### Table S1. Volatile organic compounds produced at 72h

| Chemical class | VOC name | *C. sporogenes* (n = 3) Aerobic | Anaerobic | p-value[a] | *R. lituseburensis* (n = 3) Aerobic | Anaerobic | p-value[a] | *C. subterminale* (n = 3) Aerobic | Anaerobic | p-value[a] |
|---|---|---|---|---|---|---|---|---|---|---|
| **Alkanes** | 3-methyl-pentane | - | - | - | - | - | - | 38611 ± 39965 | - | .17 |
| | Methyl-cyclopentane | - | - | - | - | - | - | 50614 ± 40103 | 30516 ± 51375 | .62 |
| | Hexane | - | - | - | - | - | - | 65701 ± 38772 | - | .04 |
| | 2,4-dimethyl-heptane | 24562 ± 22092 | 34968 ± 8798 | .49 | - | 24313 ± 42112 | .37 | 34659 ± 15258 | 306893 ± 497433 | .40 |
| | 2,2,7,7-tetramethyloctane | 40748 ± 40162 | - | .15 | - | - | - | 68252 ± 20558 | - | .004[b] |
| | 2,2,4,4-tetramethyloctane | - | - | - | 22880 ± 5845 | - | .002[b] | - | - | - |
| | 3-methyl-5-propylnonane | - | - | - | - | - | - | 25878 ± 4404 | - | .0005[b] |
| | 2,2,3-trimethylnonane | 28524 ± 28223 | - | .15 | - | - | - | 76679 ± 7809 | - | <.0001[b] |
| | 3,4-dimethyl decane | 29304 ± 26506 | - | .13 | - | - | - | 57161 ± 2709 | - | <.0001[b] |
| | 2,3,8-trimethyl-decane | 59213 ± 51582 | - | .12 | - | - | - | - | - | - |
| **Alkenes** | 1,3,5-cycloheptatriene | - | - | - | 86646 ± 58990 | - | .06 | - | - | - |
| **Aromatic compounds** | Ethylbenzene | 39496 ± 5920 | - | .0003[b] | 56380 ± 13239 | 544 ± 942 | .002[b] | 20948 ± 36283 | 6738 ± 7876 | .54 |
| | o/m/p-xylene | 19051 ± 16635 | - | .12 | 90560 ± 83629 | 735 ± 1273 | .13 | - | 15049 ± 14583 | .15 |
| **Alcohols** | 2-methylpropan-1-ol | 99224 ± 11924 | 144026 ± 28157 | .06 | - | - | - | 41237 ± 37478 | 79551 ± 22786 | .20 |
| | Butan-1-ol | - | - | - | - | - | - | 145633 ± 27551 | 290230 ± 8434 | .001[b] |
| | 2-methylbutan-1-ol | - | - | - | - | - | - | - | 59444 ± 13613 | .002[b] |
| | 3-methylbutan-1-ol | - | - | - | - | - | - | - | 92939 ± 11634 | .0002[b] |
| | 4-methylpentan-1-ol | - | 75572 ± 36979 | .02 | - | - | - | - | - | - |
| | 5-methylhexan-1-ol | - | 40101 ± 15761 | .01 | - | - | - | - | - | - |
| **SCFAE** | Butyl 2-methylpropanoate | - | - | - | - | - | - | - | 28993 ± 26891 | .14 |
| | 3-methylbutyl butanoate | - | - | - | - | - | - | - | 192737 ± 113370 | .04 |
| | Butyl 2-methylbutanoate | - | - | - | - | - | - | - | 157397 ± 94865 | .05 |
| | Butyl butanoate | - | - | - | - | - | - | - | 129407 ± 55734 | .02 |
| | 3-methylbutyl 3-methylbutanoate | - | - | - | - | - | - | 65062 ± 17208 | 808242 ± 455893 | .05 |
| | 3-methylbutyl 2-methylbutanoate | - | 273352 ± 237234 | .12 | - | - | - | - | 328103 ± 111215 | .007[b] |
| | Butyl pentanoate | - | 31786 ± 11275 | .01 | - | - | - | - | 368571 ± 236570 | .05 |
| | 2-methylpropyl pentanoate | - | 88568 ± 29681 | .01 | - | - | - | - | 111790 ± 58114 | .03 |

| | | | | | | | | | | |
|---|---|---|---|---|---|---|---|---|---|---|
| | Pentyl 4-methylpentanoate | - | 175530 ± 45957 | .003[b] | - | - | - | - | - | - |
| | Butyl hexanoate | - | 344367 ± 85369 | .002[b] | - | - | - | - | - | - |
| | 2-methylpropyl hexanoate | - | 110340 ± 34307 | .01 | - | - | - | - | - | - |
| **Sulfur compounds** | (methyldisulfanyl)methane | - | 82487 ± 4922 | <.0001[b] | 56925 ± 5420 | 201368 ± 33990 | .002[b] | - | - | - |
| | Methanethiol | - | 1120769 ± 31394 | <.0001[b] | - | 1512024 ± 141742 | <.0001[b] | - | 268622 ± 10669 | <.0001[b] |
| **Sulfur alcohols** | 2-ethenylsulfanylethanol | - | - | - | - | 116151 ± 33516 | .0001[b] | - | - | - |
| **Sulfur SCFAE** | S-methyl ethanethioate | - | - | - | - | 179067 ± 16238 | .0001[b] | - | 132276 ± 4218 | <.0001[b] |
| | S-methyl propanethioate | - | - | - | - | 48537 ± 12347 | .0001[b] | - | - | - |
| | S-methyl 2-methylpropanethioate | - | - | - | - | - | - | - | 47049 ± 3235 | <.0001[b] |
| | S-methyl butanethioate | - | - | - | - | - | - | - | 85715 ± 11373 | <.0001[b] |

Unit: peak area, -: not detected, SCFAE: Short-chain fatty acid esters. [a]two-sided t-test, [b]Discovery determined using the Two-stage linear step-up procedure of Benjamini, Krieger and Yekutieli, with Q = 1%. Each row was analyzed individually, without assuming a consistent SD.

**Table S2. Short chain fatty acids poduced at 72h**

| SCFA name | *C. sporogenes* (n = 4) | | | *R. lituseburensis* (n = 4) | | | *C. subterminale* (n = 4) | | |
|---|---|---|---|---|---|---|---|---|---|
| | Aerobiosis with antioxidants | Anaerobiosis | p-value[a] | Aerobiosis with antioxidants | Anaerobiosis | p-value[a] | Aerobiosis with antioxidants | Anaerobiosis | p-value[a] |
| Acetic acid | >10 | >10 | - | >10 | >10 | - | >10 | >10 | - |
| Propanoïc acid | 0.67 ± 0.20 | 0.84 ± 0.15 | .23 | 3.60 ± 0.56 | 1.24 ± 0.20 | .0005[b] | 0.22 ± 0.05 | 0.37 ± 0.06 | .01[b] |
| Butanoïc acid | 2.03 ± 1.04 | 1.61 ± 0.31 | .47 | 4.47 ± 0.57 | 2.93 ± 0.34 | .01[b] | 8.71 ± 0.99 | 7.26 ± 0.36 | .03[b] |
| Isobutanoïc acid | 4.05 ± 1.11 | 3.51 ± 0.72 | .44 | 4.68 ± 1.06 | 2.90 ± 0.45 | <.0001[b] | 4.68 ± 1.06 | 2.90 ± 0.45 | .02[b] |
| Isopentanoïc acid | 1.94 ± 0.31 | 2.17 ± 0.43 | .42 | 4.34 ± 0.25 | 1.28 ± 0.07 | <.0001[b] | 7.46 ± 1.59 | 7.12 ± 0.54 | .70 |

Unit: mM, Acetic acid was detected in all groups in a concentration superior to the upper limit of detection (>10mM), Pentanoïc, hexanoic, isohexanoïc and heptanoic were detected in trace amounts. All SCFA were detected in trace amounts in aerobiosis without antioxidants consistent to the absence of viability of the bacteria in this culture condition. [a]two-sided t-test, [b]Discovery determined using the Two-stage linear step-up procedure of Benjamini, Krieger and Yekutieli, with Q = 1%. Each row was analyzed individually, without assuming a consistent SD.

**Table S3. Polar metabolites detected only in aerobiosis**

| Item |
| --- |
| 3'-Uridylic acid (+) |
| 4-hydroxy-1-pyrroline-2-carboxylic acid (+) |
| Glutathione oxidised (+/-) |
| LysoPE(18:1n7/0:0) (+) |

The detection of glutathione oxidized confirmed the antioxidant activity of the glutathione included in the culture medium and is not expected to have been produced by the bacteria.

**Table S4. Polar metabolites detected only in anaerobiosis**

| Item |
|---|
| 1,4'-Bipiperidine-1'-carboxylic acid (+) |
| 3-Hydroxy-8'-apo-e-caroten-8'-al (+) |
| Cyclo(L-Trp L-Pro) (+) |
| Heptanal, 2-benzylidene- (E) (+) |
| Leucine Proline dipeptide (-) |
| Meconin (-) |
| 2-Succinyl-6-hydroxy-2,4-cyclohexadiene-1-carboxylate (SHCHC) (-) |

**Figure S1. Volatile organic compounds produced according to time**

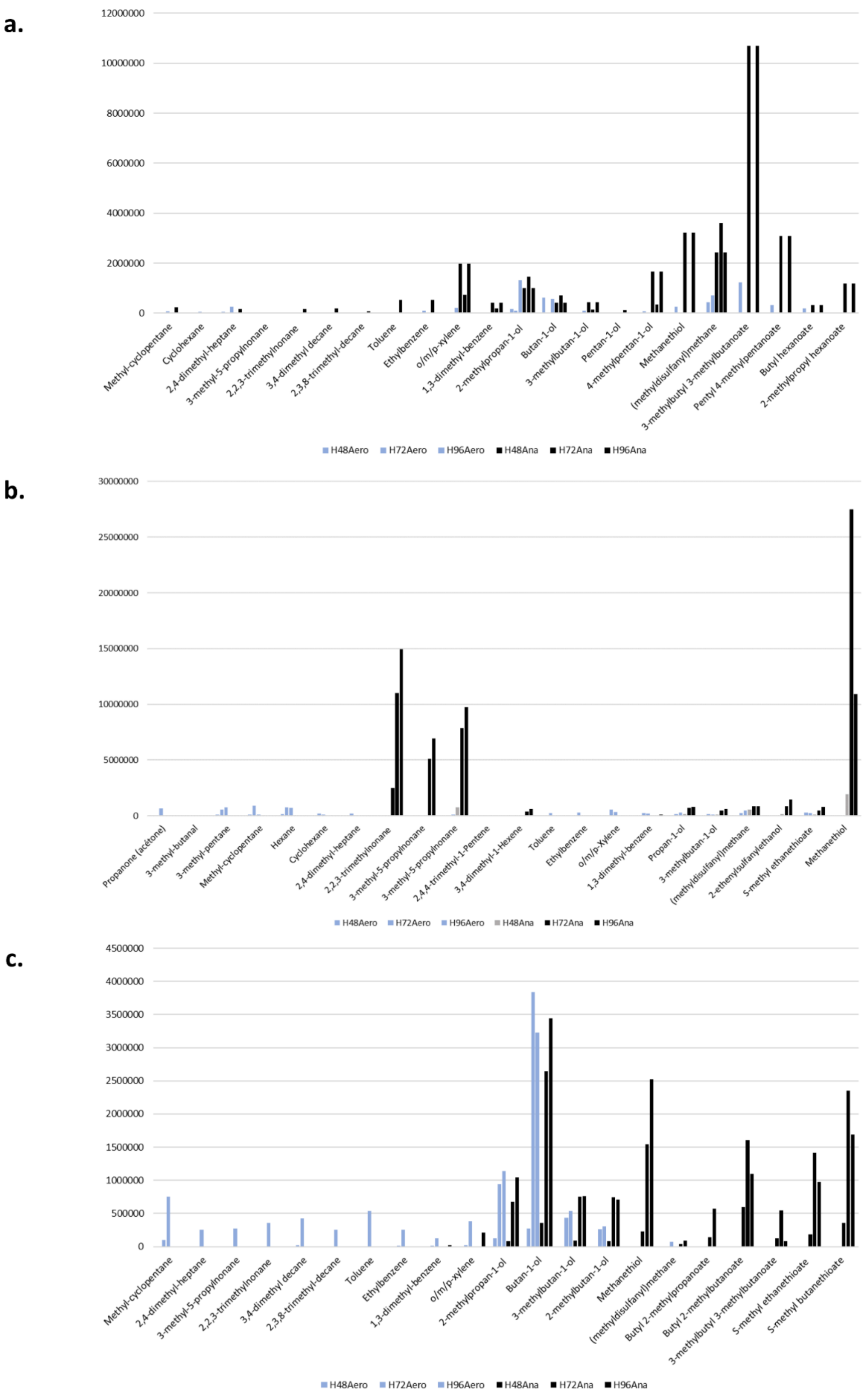


a. *Clostridium sprogenes*, b. *Romboutsia lituseburensis*, c. *Clostridium subterminale*. Arbitrary unit (area under peak).

**Figure S2. Phylogenetic tree of the three strains used in this study**

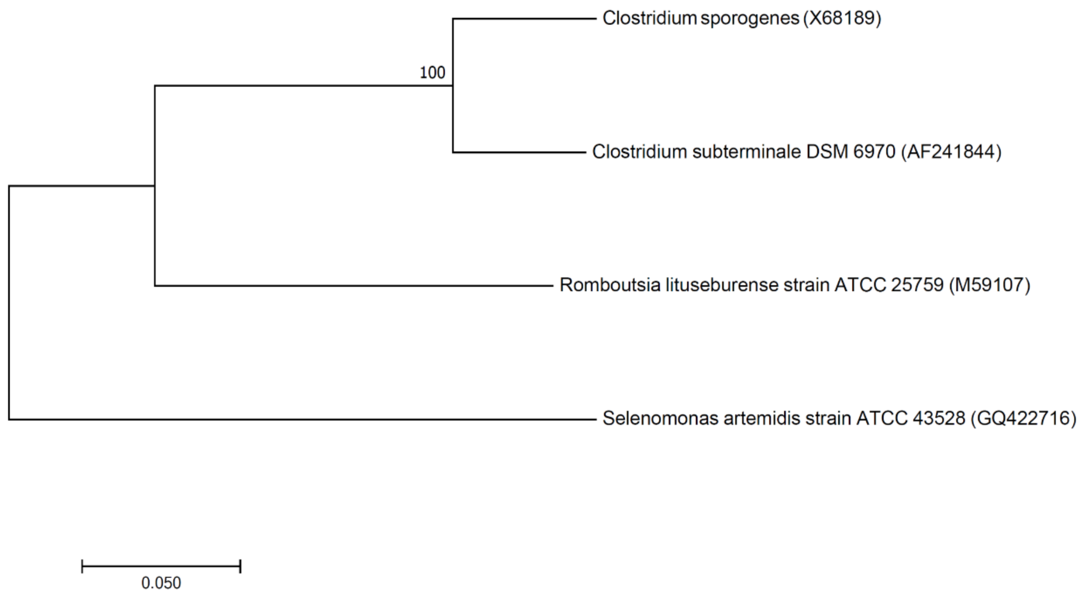


For each of the 3 species, sequence accession no. (16S rRNA gene) for the type strain was obtained from the taxonomic reference site http://www.bacterio.net/ for each species. The Genbank sequence number were as follows: *Clostridium sporogenes*: X68189, *Clostridium subterminale*: AF241844, *Romboutsia lituseburensis* : M59107. Consistently with the nomenclature, *C. sporogenes* and *C. subterminale* are more closely related than with *R. lituseburensis*. The tree was performed with the Molecular Evolutionary Genetic Analysis Software MEGA v7.0.26 (https://www.megasoftware.net/).[1]

**Figure S3. Volatile organic compounds produced by each species in aerobiosis with antioxidants or in anaerobiosis**

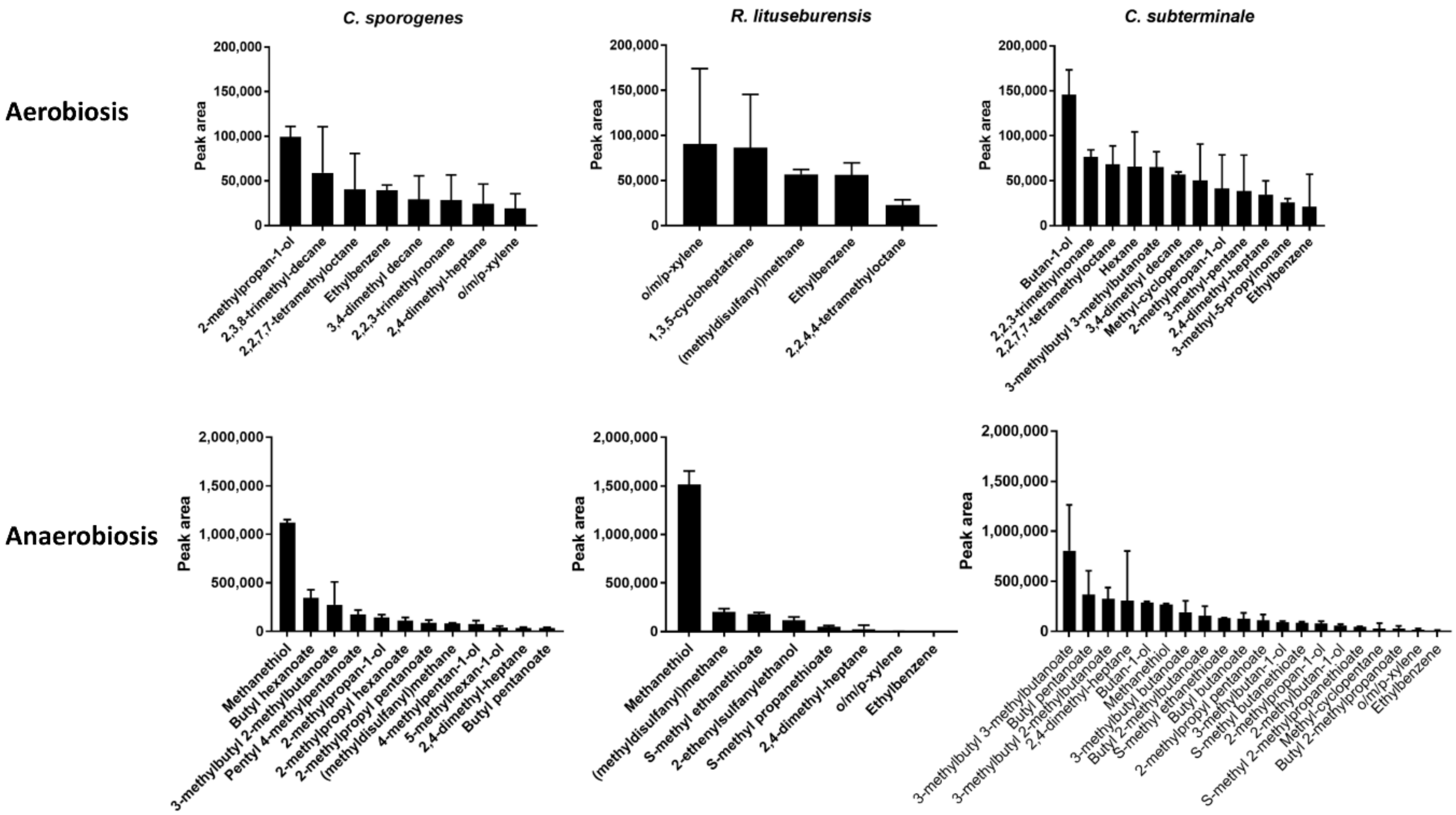

**Figure S4. Biplot of odors according to species and atmosphere**

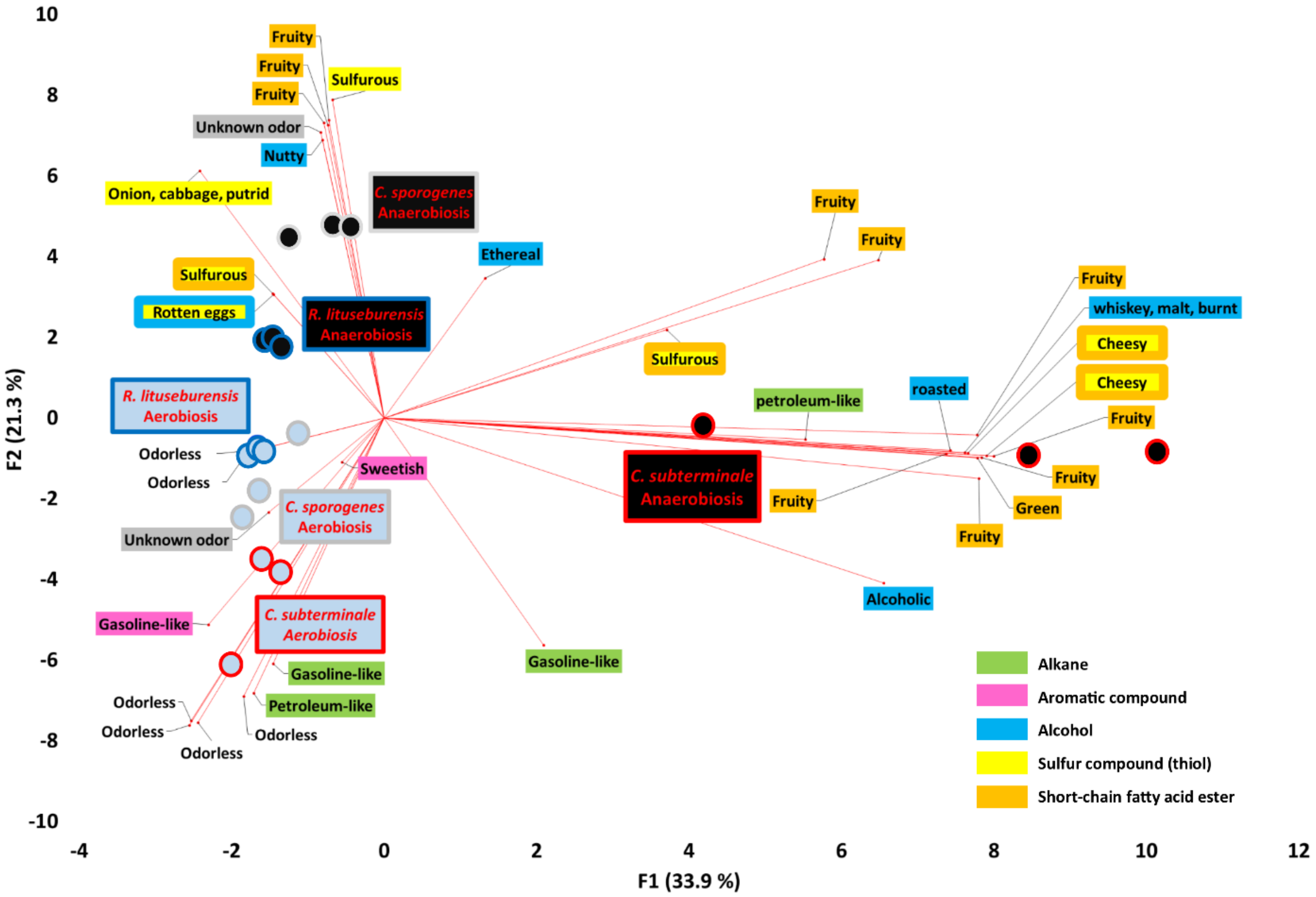
